# Relationships between macro-fungal dark diversity and habitat parameters using LiDAR

**DOI:** 10.1101/2020.07.02.185553

**Authors:** Jose W. Valdez, Ane Kirstine Brunbjerg, Camilla Fløjgaard, Lars Dalby, Kevin K. Clausen, Meelis Pärtel, Norbert Pfeifer, Markus Hollaus, Michael H. Wimmer, Rasmus Ejrnæs, Jesper Erenskjold Moeslund

## Abstract

Despite the important role of fungi for ecosystems, relatively little is known about the factors underlying the dynamics of their diversity. Moreover, studies do not typically consider their dark diversity: the species absent from an otherwise suitable site. Here, we examined potential drivers of local fungal dark diversity in temperate woodland and open habitats using LiDAR and in-situ field measurements, combined with a systematically collected and geographically comprehensive macro-fungi and plant data set. For the first time, we also estimated species pools of fungi by considering both plant and fungi co-occurrences. The most important LiDAR variables for modelling fungal dark diversity were amplitude and echo ratio, which are both thought to represent vegetation structure. These results suggest that the local fungal dark diversity is highest in production forests like plantations and lowest in more open forests and in open habitats with little woody vegetation. Plant species richness was the strongest explanatory factor overall and negatively correlated with local fungal dark diversity. Soil fertility showed a positive relationship with dark diversity in open habitats. These findings may indicate that the local dark diversity of macro-fungi is highest in areas with a relatively high human impact (typically areas with low plant species richness and high soil fertility). Overall, this study brings novel insights into local macro-fungi dark diversity patterns, suggesting that a multitude of drivers related to both soil and vegetation act in concert to determine fungal dark diversity.

## Introduction

Understanding the underlying drivers shaping biodiversity patterns is a central goal in ecology and conservation biology. This is also true for fungi which play a vital role in ecosystem functioning as decomposers, mutualists, and pathogens. However, fungi and the underlying environmental factors influencing fungal diversity is less studied than animals and plants, and quantifying fungal diversity is far from trivial. The most commonly used biodiversity metric is observed species richness (Mueller 2011). However, this measure is not always suitable for comparisons across habitats and conveys no information on the part of the diversity that is potentially missing in a given site (Pärtel *et al.* 2011). In addition, monitoring fungal diversity is often severely hampered by detectability issues and the life history of the involved species (Yahr *et al.* 2016, Blackwell and Vega 2018). Several alternative approaches have been developed to more effectively monitor and compare biodiversity across landscapes (Solow and Polasky 1994, Sarkar and Margules 2002, Ricotta 2005, Ricotta 2007). Although these methods can provide valuable insights, they do not consider the dark diversity, the absent part of the species pool which can potentially inhabit an environmentally suitable site (Pärtel *et al.* 2011). This often-ignored aspect of diversity provides a novel and ecologically meaningful metric for estimating how much of the potential species diversity – the site-specific species pool – is lacking (Pärtel *et al.* 2011). This information is important to understand the underlying mechanisms and dynamics of community assembly (e.g., community saturation) (Mateo *et al.* 2017). Dark diversity may also become an important conservation tool to measure biodiversity potential, such as aiding managers or policy-makers to prioritize certain habitats, estimate restoration potential of degraded habitats, or forecast potential impacts of invasions (Lewis *et al.* 2017). Here, we use fungal data from 130 thoroughly inventoried sites covering all terrestrial habitats, from open to forest, and wet to arid, to investigate important drivers of fungal dark diversity.

Dark diversity aims to reconcile the role of simultaneous, and potentially confounding, regional and local processes underlying biodiversity patterns and biological communities (Pärtel *et al.* 2011, Pärtel 2014). In any given landscape, the biodiversity potential is ultimately determined by large-scale biogeographic and evolutionary processes (i.e., species diversification and historic migration patterns) determining the set of species which can theoretically inhabit a site, defined as the regional pool (Pärtel *et al.* 1996, Cornell and Harrison 2014, Zobel 2016). This regional pool is further filtered by local processes such as environmental gradients, species interactions, population dynamics, dispersal, disturbance, and stochastic events and referred to as the site-specific species pool, i.e., species that could possibly live in a given site (Pärtel *et al.* 2013, Cornell and Harrison 2014, Ronk *et al.* 2015, Zobel 2016). While many studies have investigated the drivers of fungal diversity, only a few studies have focused on the determinants of fungal dark diversity. These studies demonstrate that higher temperatures increases arbuscular mycorrhizal dark diversity (Pärtel *et al.* 2017a) and annual precipitation decreases the dark diversity of ectomycorrhizal fungi at the global scale (Pärtel *et al.* 2017b). These results concur with previous research suggesting that large scale climatic factors are strong drivers of fungal richness and community composition, attributed to the direct and indirect effects which alter soil and floristic conditions (Staddon *et al.* 2003, Kivlin *et al.* 2011, Tedersoo *et al.* 2014). Local edaphic conditions such as soil moisture, pH, and calcium concentration are also known to influence fungal diversity (Geml *et al.* 2014, Tedersoo *et al.* 2014, Tonn and Ibáñez 2017, Frøslev *et al.* 2019), but it is not known how these environmental factors affect fungal dark diversity. In fact, the general mechanisms determining dark diversity in fungal communities remain largely unknown. Clearly, species can be absent from an area just by chance (Hubbell 2011). Species can also be absent from a site because of some kind of unexpected disturbance – for example human, but it could also be natural – altering species’ dispersal, establishment, or persistence possibilities. In principle, these disturbances can be both biological and chemical and act at various spatial scales. An example could be extreme drought. Often habitats have a constant level of relatively low or usual disturbances that the habitat’s species are adapted to (e.g., grazing), and these do not count towards disturbances that can cause dark diversity. It is important not to confuse dark diversity with hidden diversity; i.e., species that are actually present in a given site but just not recorded (Milberg *et al.* 2008, Abrego *et al.* 2016).

Besides the influence of environmental gradients, other factors particularly important for fungi are vegetation and habitat structure, such as vegetation height, shrub layer, vegetation cover, dead wood, and other woody features (Humphrey *et al.* 2000, Nordén and Paltto 2001, Nordén *et al.* 2004, Gómez-Hernández and Williams-Linera 2011, Zuo *et al.* 2016). As the dominant primary producer in terrestrial ecosystems, plants also form the living and dead organic carbon pools and biotic surfaces that are the niche space for not only fungi but other taxonomic groups as well (DeAngelis 2012, Brunbjerg *et al.* 2017). These structural elements are an important element for biodiversity, and can influence not only fungal diversity, but the diversity of plants, animals, and bacteria as well (Penone *et al.* 2019). However, despite the obvious contribution of these variables, such factors are rarely covered extensively since they are difficult to measure and require large amounts of resources to obtain sufficient and high quality data. However, emerging technologies such as LiDAR (light detection and ranging) could potentially remedy this situation.

Airborne LiDAR records a three-dimensional set of points using laser ranging from an aircraft or a drone (Lefsky *et al.* 2002). It captures data suitable to represent many of the vegetation and landscape structural measures important to fungi (Vehmas *et al.* 2009, Lopatin *et al.* 2016, Peura *et al.* 2016, Thers *et al.* 2017, Mao *et al.* 2018). As a relatively new methodology, biodiversity studies that employ LiDAR have been limited in scope, typically addressing only one taxonomic group or habitat type at the local scale, and strongly biased towards forest ecosystems. However, studies using LiDAR-based indicators have already been shown to explain up to 66% and 82% of local plant and fungi richness, respectively (Lopatin *et al.* 2016, Peura *et al.* 2016, Thers *et al.* 2017). A recent study has demonstrated its potential to provide spatially accurate and comprehensive measures by predicting the local biodiversity of different taxonomic groups (plants, fungi, lichens, and bryophytes) across multiple habitat types and large geographic extent (Moeslund *et al.* 2019). LiDAR may also be a useful tool in studying dark diversity by incorporating potentially important spatiotemporal dynamics such as succession and disturbance (Mokany and Shine 2003, Scott *et al.* 2011, Pärtel *et al.* 2013). For example, recent studies have found that human impact increases dark diversity in arbuscular mycorrhizal fungi (Pärtel *et al.* 2017a), that ruderal plants are more likely to be in dark diversity (Moeslund *et al.* 2017), and that human density and agricultural land-use influence dark diversity of vascular plants (Riibak *et al.* 2017).

Alongside these structural and environmental factors, fungal diversity depends on biotic interactions, with a large proportion of fungi deriving their nutrients and carbon from host plants (Tedersoo *et al.* 2014, Nguyen *et al.* 2016). Recent evidence has hinted on the influence of these interactions on dark diversity, as plant species dependent on mycorrhiza have been found to have greater dark diversity than those without these mutualist relationships (Moeslund *et al.* 2017). Moreover, ectomycorrhizal fungal diversity seems to increase exponentially with an increasing proportion of their host plants, suggesting that competitive interactions among fungi might also drive their dark diversity (Pärtel *et al.* 2017b). Typical and strong species interactions are indeed typically considered in dark diversity, as the estimation hereof is usually based on species co-occurrences (Beals 1984, McCune 1994, Münzbergová and Herben 2004, de Bello *et al.* 2012, Lewis *et al.* 2016). However, this is usually done only within the species group being studied. For example, in studies of plant dark diversity, only co-occurrences with other plants, and not fungi or other species groups, are typically considered. However, recognizing the close and interconnected relationship between plants and fungi allows for stronger and more realistic estimations of the fungal dark diversity. Incorporating other taxonomic groups when determining species pools and estimating dark diversity is not a new insight, and the importance of biotic interactions across trophic groups has been discussed since the concept of dark diversity was first introduced (Pärtel *et al.* 2011). Yet, such cross-species group data has never been included in dark diversity estimates meaning that they may not sufficiently account for cross-species group interactions, and this may be part of the explanation for why dark diversity is sometimes over-estimated (Boussarie *et al.* 2018).

In this study, we examined a number of environmental factors influencing the local dark diversity of fungi across habitat-types nationwide within Denmark. We used a comprehensive biodiversity datasets covering major environmental gradients (Brunbjerg *et al.* 2019) and combined it with LiDAR-based measurements. We also included fungi-plant-co-occurrence information to estimate local fungal dark diversity and thereby acknowledge the importance of their biotic interactions. More specifically, we addressed the following questions: 1. To what degree can we explain local fungal dark diversity by abiotic and biotic environmental factors? 2. Can vegetation and terrain structural factors that are important to local fungal dark diversity be derived from LiDAR and if so, 3. how important are they compared to field-measured factors?

## Methods

### Study area and site selection

The dataset was collected from a national biodiversity inventory in Denmark as part of the “Biowide” research project (Brunbjerg *et al.* 2019). A total of 130 study sites (40 × 40 m) were selected with a minimum distance of 500 m between each to reduce spatial covariance with 30 sites allocated to cultivated habitats and 100 sites to natural habitats (Figure 1). The cultivated subset was stratified according to the type of land use and the natural subset was selected amongst uncultivated habitats and stratified according to gradients in soil fertility, soil moisture, and successional stage. The “Biowide” project deliberately excluded saline and aquatic habitats but included temporarily inundated depressions along with mires, bogs, and fens. The final set of 24 habitat strata consisted of three types of fields (rotational, grass leys, set aside) and three types of plantations (beech, oak, spruce), and the remaining 18 strata were natural or semi-natural habitats, constituting all possible combinations of positions along three major natural environmental gradients: soil fertility (rich, poor), soil moisture (dry, moist, wet), and successional stage (early, mid, late). These 24 strata were replicated in five geographical regions in Denmark. The “Biowide” dataset also includes a subset of 10 sites (two in each region) of hotspots for different taxonomic groups in Denmark, which were selected by voting amongst active naturalists in the Danish conservation and management societies. For further details on the design and data collection procedures see Brunbjerg *et al.* (2019).

**Figure 1.**
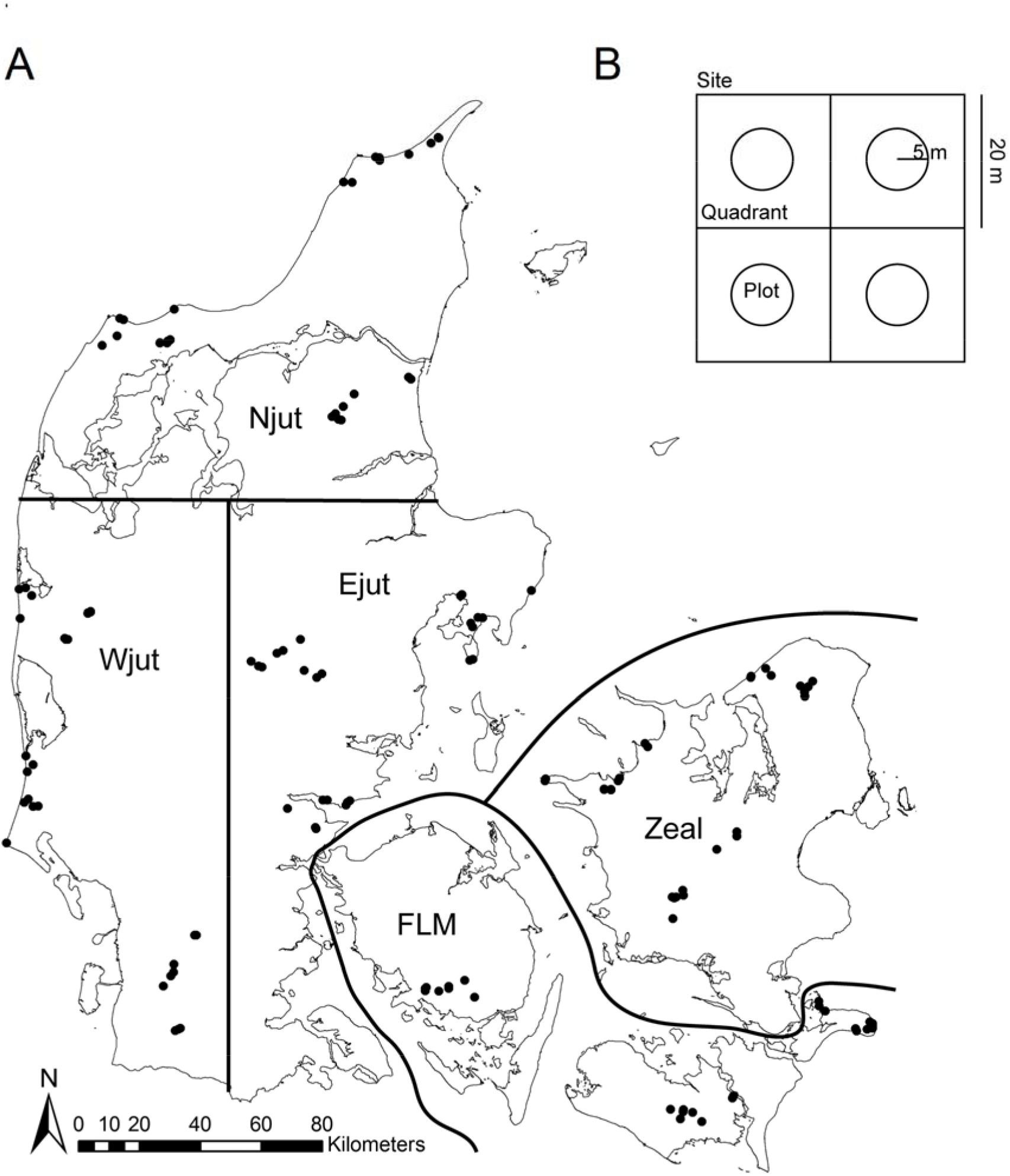
The (A) 130 selected study sites from a national biodiversity inventory and (B) the four quadrants and 5 m circle plots within each site. Reprinted from Ejrnæs *et al.* (2018), with permission from Elsevier.

### Field-measured variables

We used fungi observational data from the “Biowide” field inventories (summarized data and additional details in Brunbjerg *et al.* (2019)). Macro-fungal species were surveyed in 2014–2015 by an expert field-mycologist and volunteers during three inventories (up to one hour per site) in the main fruiting season (August - November) by actively searching microhabitats and substrates (soil, herbaceous vegetation and debris, dead wood, litter, and bark of trees up to 2 m) within the 40 × 40 m sites. Since truffles are difficult to find, we did not consider these in this study. Subspecies and varieties were lumped to the species level. After pre-processing, the dataset consisted of 6,269 observations of 1,017 species.

Vascular plant species observations were also taken from the “Biowide” database and were originally inventoried by trained botanists during the summer 2014 and spring 2015 to account for variations in phenology. We removed hybrids and neophytes (i.e. species that are not considered a natural part of the vegetation given their history and dispersal ability, see appendix tables 6–8 in Buchwald *et al.* (2013)) and lumped subspecies and varieties to the species level. Species nomenclature for both plants and fungi follow the species checklist of Denmark (allearter.dk).

Apart from the LiDAR-based measures (detailed below), we also considered field-measured variables representing both abiotic conditions and available biotic resources known to influence fungal diversity and communities (Table 1). For further details on data collection and how the environmental field measurements were made see Brunbjerg *et al.* (2019).

**Table 1.**
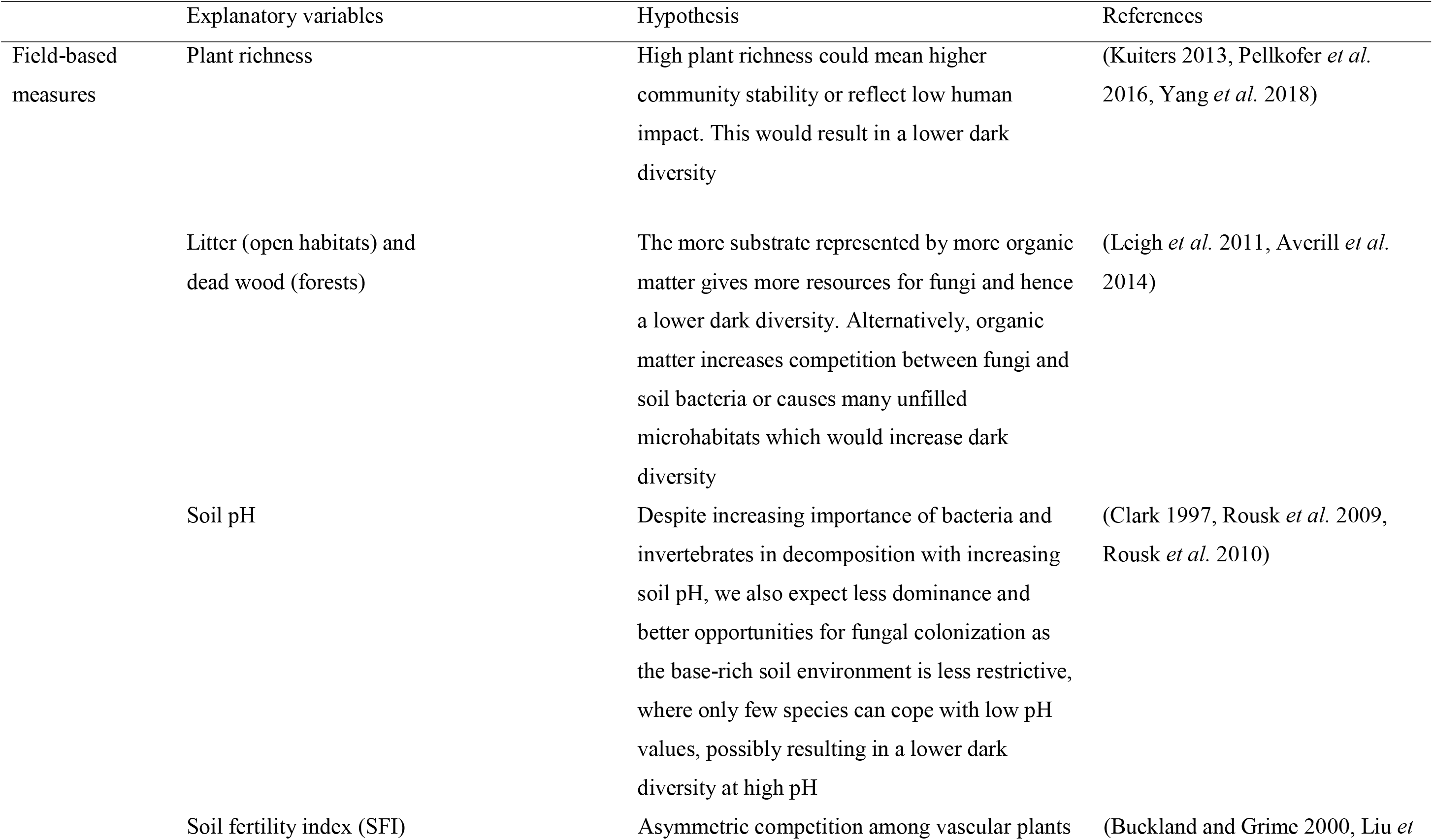
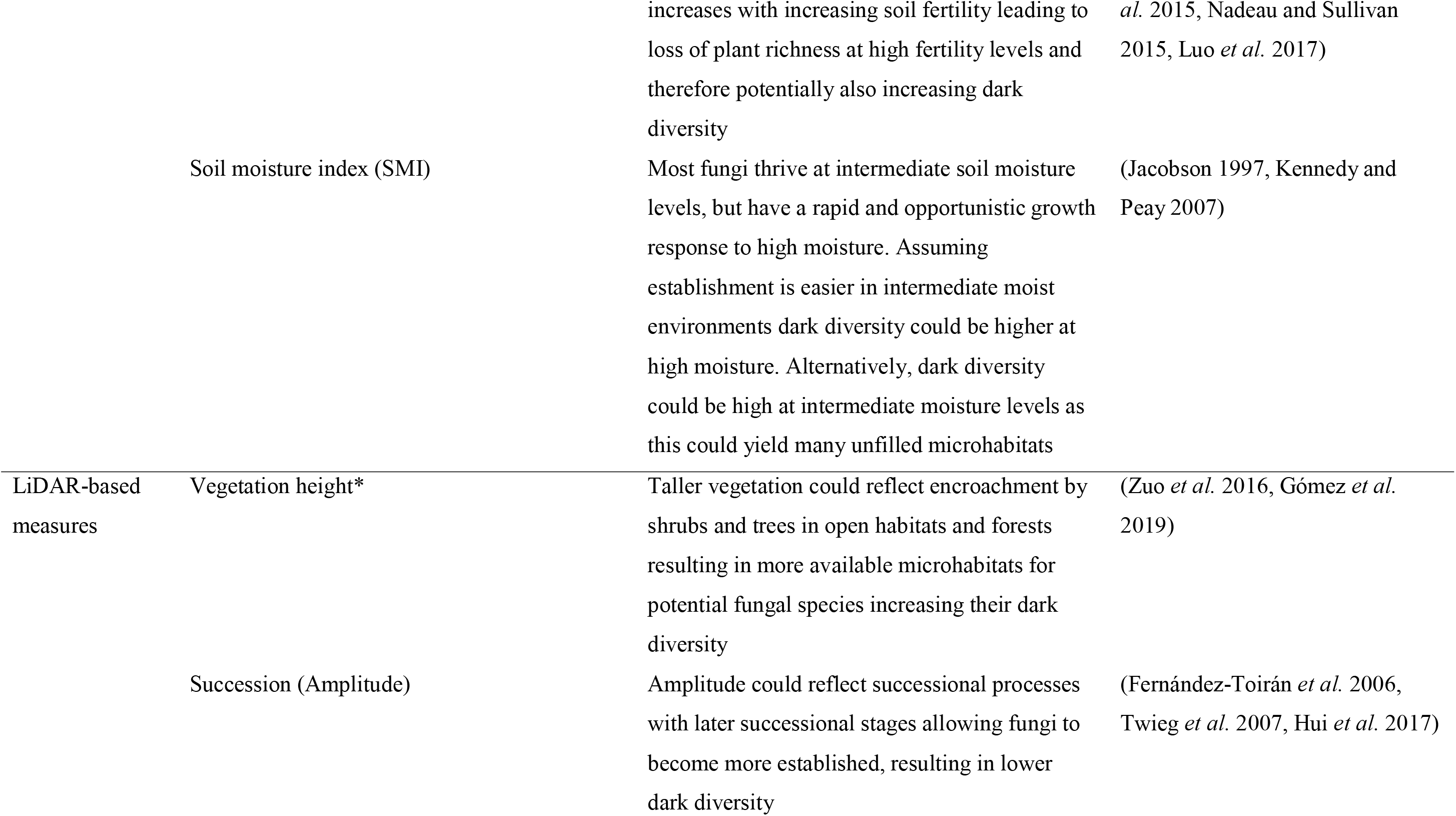
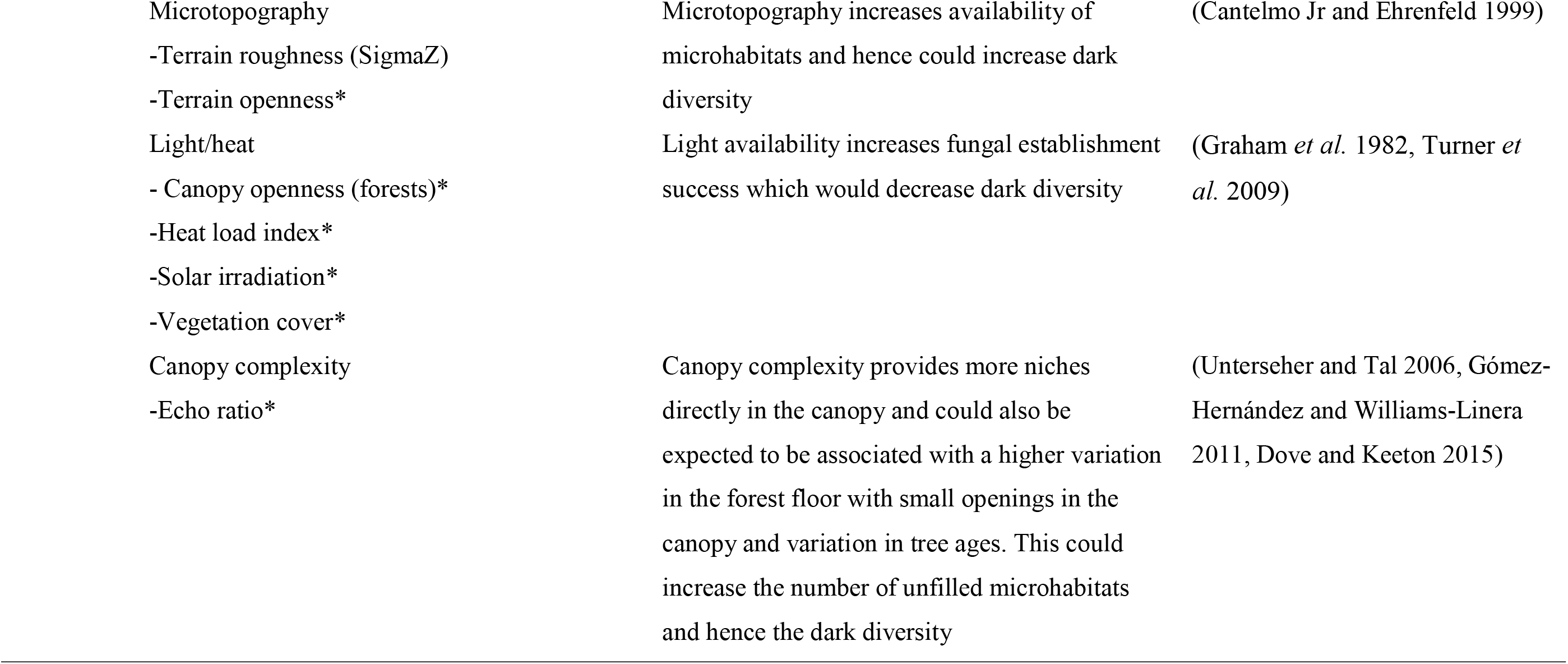
Overview of the explanatory variables for fungal dark diversity models along with our hypothesized relationship with dark diversity. If the standard deviation of a variable was calculated, in addition to its mean, the variable is denoted with an asterisk.

|  | Explanatory variables | Hypothesis | References |
| --- | --- | --- | --- |
| Field-based measures | Plant richness | High plant richness could mean higher community stability or reflect low human impact. This would result in a lower dark diversity | (Kuiters 2013, Pellkofer *et al.* 2016, Yang *et al.* 2018) |
|  | Litter (open habitats) and dead wood (forests) | The more substrate represented by more organic matter gives more resources for fungi and hence a lower dark diversity. Alternatively, organic matter increases competition between fungi and soil bacteria or causes many unfilled microhabitats which would increase dark diversity | (Leigh *et al.* 2011, Averill *et al.* 2014) |
|  | Soil pH | Despite increasing importance of bacteria and invertebrates in decomposition with increasing soil pH, we also expect less dominance and better opportunities for fungal colonization as the base-rich soil environment is less restrictive, where only few species can cope with low pH values, possibly resulting in a lower dark diversity at high pH | (Clark 1997, Rousk *et al.* 2009, Rousk *et al.* 2010) |
|  | Soil fertility index (SFI) | Asymmetric competition among vascular plants | (Buckland and Grime 2000, Liu *et* |
|  |  | increases with increasing soil fertility leading to loss of plant richness at high fertility levels and therefore potentially also increasing dark diversity | *al.* 2015, Nadeau and Sullivan 2015, Luo *et al.* 2017) |
|  | Soil moisture index (SMI) | Most fungi thrive at intermediate soil moisture levels, but have a rapid and opportunistic growth response to high moisture. Assuming establishment is easier in intermediate moist environments dark diversity could be higher at high moisture. Alternatively, dark diversity could be high at intermediate moisture levels as this could yield many unfilled microhabitats | (Jacobson 1997, Kennedy and Peay 2007) |
| LiDAR-based measures | Vegetation height* | Taller vegetation could reflect encroachment by shrubs and trees in open habitats and forests resulting in more available microhabitats for potential fungal species increasing their dark diversity | (Zuo *et al.* 2016, Gómez *et al.* 2019) |
|  | Succession (Amplitude) | Amplitude could reflect successional processes with later successional stages allowing fungi to become more established, resulting in lower dark diversity | (Fernández-Toirán *et al.* 2006, Twieg *et al.* 2007, Hui *et al.* 2017) |

|  |  |  |
| --- | --- | --- |
| Microtopography | Microtopography increases availability of | (Cantelmo Jr and Ehrenfeld 1999) |
| -Terrain roughness (SigmaZ) | microhabitats and hence could increase dark |  |
| -Terrain openness* | diversity |  |
| Light/heat | Light availability increases fungal establishment | (Graham *et al.* 1982, Turner *et* |
| - Canopy openness (forests)* | success which would decrease dark diversity | *al.* 2009) |
| -Heat load index* |  |  |
| -Solar irradiation* |  |  |
| -Vegetation cover* |  |  |
| Canopy complexity | Canopy complexity provides more niches | (Unterseher and Tal 2006, Gómez- |
| -Echo ratio* | directly in the canopy and could also be | Hernández and Williams-Linera |
|  | expected to be associated with a higher variation | 2011, Dove and Keeton 2015) |
|  | in the forest floor with small openings in the |  |
|  | canopy and variation in tree ages. This could |  |
|  | increase the number of unfilled microhabitats |  |
|  | and hence the dark diversity |  |

### LiDAR-based measures

To enable the calculation of measures representing vegetation and terrain environmental and structural aspects, we used the latest nationally covering LiDAR-based point cloud for Denmark from the Danish Ministry of Environment. This dataset is freely available from www.kortforsyningen.dk and has a point density of 4-5 points/m^2^. Originally, this dataset was recorded from fixed-wing airplanes at an altitude of approximately 680 m above ground level and a speed over ground of approximately 240 km/h. The data was recorded by Riegl LMS-680i scanners operating in the near-infrared wavelength (1550 nm) in a parallel line scan pattern during the springs and autumns of 2014 and 2015. For all calculations, we relied on the classification of points into ground, building and vegetation classes already present in the data set upon download.

To represent vegetation and terrain environmental and structural aspects, we calculated observed measures based on the point cloud data set. We calculated all measures at 1.5 m resolution (except for terrain roughness which was at 0.5 m resolution) and their means and standard deviations within 30 m radius circles centered in each study site. For all LiDAR processing and calculation, we used the OPALS tools (Pfeifer *et al.* 2014) version 2.3.1 in a Python 2.7 environment.

#### Vegetation-related measures

To represent *succession* and to some degree moisture balance in both vegetation and soil, we used the amplitude of each echo representing a point in the LiDAR point cloud. This amplitude is high if the reflecting surface is flat (i.e., smooth) and with high reflectivity. It is low when the light energy is distributed between several returns for example in tree canopies, or when surfaces have low reflectivity, are complex, or translucent (e.g., leaves). The wavelength used to record the point cloud data is sensitive to leaf water content (Junttila *et al.* 2018) and soil moisture (Zlinszky *et al.* 2014). Since the amplitude depends on reflectivity, which varies across months and aircraft types (slightly different flying heights) used for data recording, the amplitude was corrected to account for these biases. We constructed a Generalized Linear Model (GLM) with Gaussian link having the raw amplitude as response and flight month as well as aircraft type as explanatory factors and used only the residuals of this model for input in our statistical modelling. We also tried using flight year as an explanatory factor, but this did not improve the model (ΔAIC < 2). These residuals will be referred to as the *corrected amplitude* in the following. Unfortunately, we did not have reference data enabling a full calibration of this measure (Höfle and Pfeifer 2007).

To represent *vegetation height*, we estimated this measure by subtracting the terrain model from the surface model (two raster files, detailed in the following). The terrain model (DTM) calculation details are given in the section on “Terrain-structure measure”. The surface model was calculated using the DSM module in OPALS using all vegetation and ground points.

To reflect the penetrability and succession of the vegetation we calculated the *echo ratio* (Höfle *et al.* 2012). Echo ratio is high where the surface is impenetrable and relatively smooth and lower where the surface is uneven or penetrable. In order to calculate the echo ratio, estimating normals for each point is required. We did this using the Normals module in OPALS with a robust plane fit based on the 12 nearest neighboring points. Subsequently, we calculated the echo ratio for each terrain and vegetation point using a search radius of 1.5 m along with the slope adaptive method implemented in the EchoRatio module of OPALS.

To estimate light conditions, we calculated the *canopy openness* for all points categorized as “ground”, but contrary to terrain openness (see below), we calculated this considering vegetation points as well. Therefore, canopy openness represents the actual blocking of the sky view by the canopy around each ground point. Canopy openness is high for ground points inside canopy gaps and low for ground points beneath a closed canopy.

Lastly, as an estimate of *vegetation cover*, we calculated the fraction of vegetation points to all points (excluding unclassified points and those representing buildings and noise). This measure will be high if the vegetation is dense or the cover of vegetation is relatively high, and low for areas with no vegetation.

#### Terrain-structure measures

To enable the calculation of several terrain-related measures, we calculated a digital terrain model (DTM) for each study site representing the elevation above sea level. To do this we used the DTM module of OPALS based on only ground points. We set the module to use 8 neighboring points and a search radius of 6 m. To represent key features of the local terrain (e.g., soil moisture or heat balance (Moeslund *et al.* 2013)), we calculated *terrain slope* and *terrain aspect* (used for heat load index calculation, see below). For this task, we used the GridFeature module of OPALS using the DTM as input, a kernel size of 1 and requesting the terrain slope and aspect (slope direction) in radians.

To reflect local heat input, we calculated the *heat load index* based on the terrain aspect following the heat load index formula in McCune and Keon (2002). This index reaches maximum values on southwest-facing slopes and zero on northeast-facing slopes. We also calculated the potential *solar irradiation* based on terrain slope, aspect, and latitude following equation 3 in McCune and Keon (2002).

To estimate micro-scale terrain heterogeneity, we calculated the *terrain roughness* (SigmaZ) using only ground points as input. This measure represents the standard deviation of the interpolated grid height. The OPALS DTM module outputs this measure as a by-product when constructing a DTM. However, unlike the rest of the LiDAR measures in this study, the terrain roughness was calculated at 0.5 × 0.5 m resolution mirroring micro-scale terrain variations.

To represent site-scale terrain heterogeneity, we calculated the *terrain openness* (Doneus 2013). Terrain openness is defined as the opening angle of a cone (having the radius of the kernel) turned upside down – with its tip restrained to the point of interest – that touches the terrain surface. To calculate this, we used the PointStats module of OPALS requesting “positive openness” based on only ground points and a search radius of 5 m. This measure is high in flat (relative to the scale at which it is calculated) areas and low in heterogeneous terrains.

Finally, to test the importance of variability in the LiDAR measures we calculated the standard deviation for LiDAR measures for which we believed it made ecological sense (Table 1).

### Data analysis

#### Data preparation

Prior to statistical analysis, we removed the six intensively managed fields from the study sites, as these were ploughed fields. We also removed two study sites because they were flooded during the LiDAR data recording period. Finally, we removed one site due to an extreme outlier in the LiDAR amplitude values (300 vs. a range of values between 10 and 130). Our final dataset therefore comprised a total of 121 study sites.

Our initial visual inspection of the data revealed that many of the LiDAR measures were relevant only for woodlands and therefore strongly zero-inflated in the open landscapes. The analyses in this study were therefore separately run for open habitats and woodlands. Open habitats included grasslands, fens, bogs, and other habitats with only few sporadic occurrences of trees. The woodlands dataset consisted of forests, thickets and shrubland (e.g. willow).

In the following, we detail the steps we took to prepare the LiDAR and measured variables for statistical modelling as explanatory factors. Obviously, a number of these variables were strongly inter-correlated (Appendix 1). For example, echo ratio was strongly related to canopy and light measures (Appendix 1). Therefore, we selected only those variables that we hypothesized to affect local fungal dark diversity (See Table 1). Subsequently, to avoid issues with multi-collinearity we calculated Variance Inflation Factors (VIFs) causing us to remove vegetation height as an explanatory factor for the open landscapes to ensure VIF values below 10 following Kutner *et al.* (2005). Subsequently, the maximum VIF value of explanatory factors used together in the same models was 4.8 and 5.7 for woodlands and open habitats respectively. We scaled all explanatory variables to a mean of zero and a standard deviation of one. To strive for normal distribution of explanatory variables we log- or square-root transformed those where this made obvious distributional improvements based on visual examination of the histograms.

**Appendix 1.**
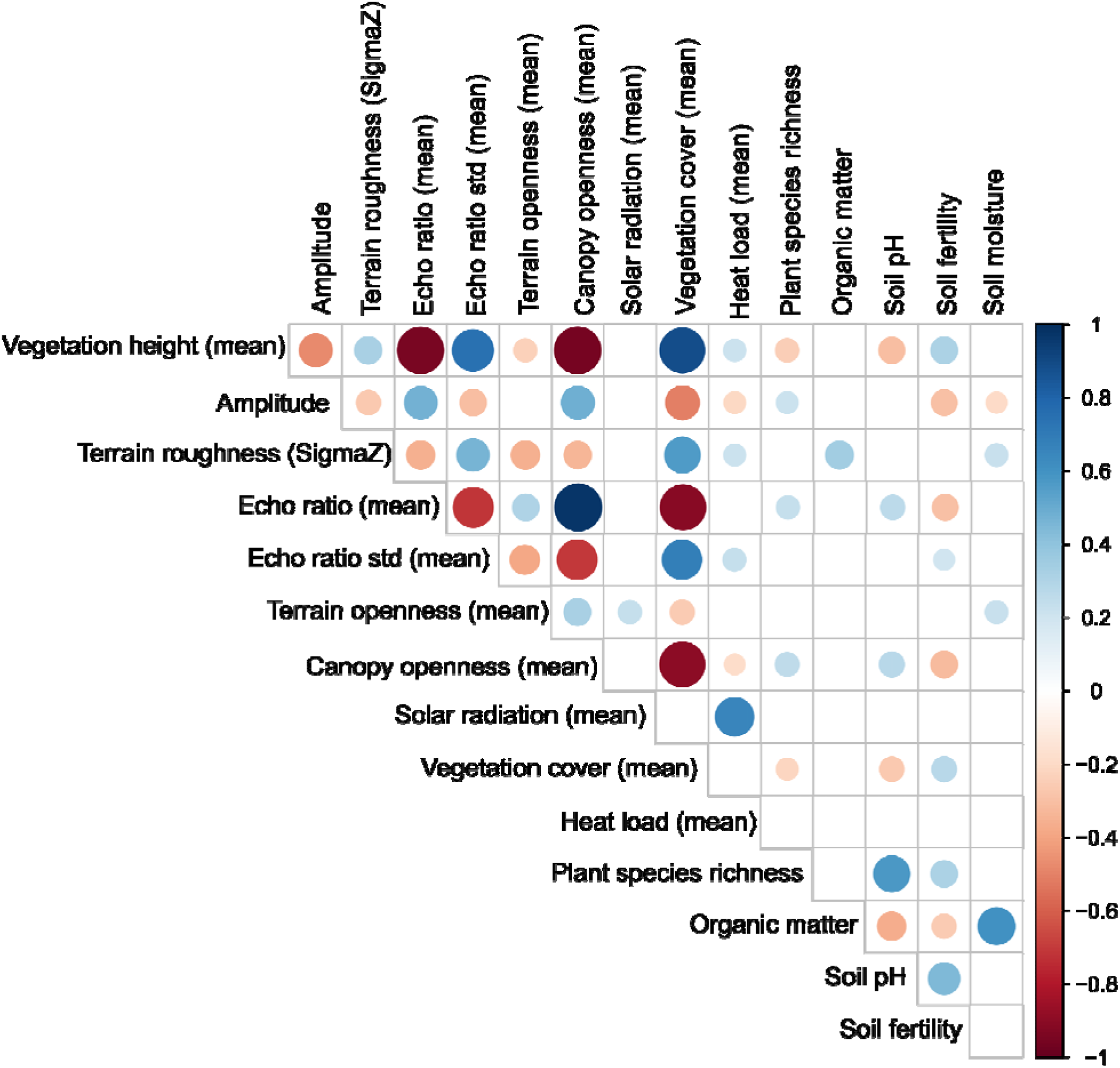
Correlation plot between environmental and LiDAR variables. All significant interactions are colored red for negative relationships and blue for positive relationships, with the size and darkness of the color representing the strength of the relationship. Non-significant correlations are blank.

#### Dark diversity

All statistical analyses mentioned in the following were performed in R version 3.5.3 (**R** Core Team 2019). To calculate the dark diversity for each site, we first calculated the site-specific species pool using the Beals’ index (Beals 1984), as recommended by Lewis *et al.* (2016), using the ‘beals’ function in the ‘vegan’ package (Oksanen *et al.* 2017). The Beals’ index represents the probability that a particular species will occur within a given site based on the assemblage of co-occurring species (Beals 1984, McCune 1994, Münzbergová and Herben 2004). The threshold used for including a particular species in the site-specific species pool was the 5^th^ percentile of the Beals’ index value for each species (Gijbels *et al.* 2012, Ronk *et al.* 2015). Preceding the calculation of each threshold, the lowest Beals’ index value among plots with the occurrence of the species in question was identified, and all plots having values below the minimum were not considered. We calculated two measures of the site-specific species pool for each site: (1) using only fungi co-occurrence and (2) co-occurrences of both observed fungi and vascular plants at each site to acknowledge the fungal-plant linkages (i.e., both fungi and plant species were in the presence/absence matrix used to calculate Beals’ index). Dark diversity was calculated by subtracting observed fungal species richness from the site-specific species pool. Since site-specific species pools differ between sites, we calculated the *relative dark diversity* for each site as dark diversity (species predicted from the site-specific species pool but not observed) divided by the regional pool to enable comparison of results across habitats.

#### Statistical analysis

To investigate what characterizes sites with a high fungal dark diversity we constructed GLMs with a Gaussian link having the estimated relative dark diversity (described above) as the response variable. We constructed models for both open habitats and woodlands, and for both dark diversity estimates (see the section on dark diversity). Initially, we fitted models using only the LiDAR measures as explanatory factors, to test the degree to which fungal dark diversity patterns could be explained using LiDAR data alone. Subsequently, we fitted a similar model with both measured and LiDAR variables as explanatory factors (Table 1), giving insight into how much more explanatory power one gains by using measured variables in addition to LiDAR. To allow for non-linear relationships for variables corresponding to the intermediate disturbance hypothesis (Connell 1978, Townsend *et al.* 1997) and intermediate productivity hypothesis (Fraser *et al.* 2015), we used Akaike’s Information Criterion (AIC) (Burnham and Anderson 2002) to evaluate if inclusion of squared terms for the variables SMI, SFI, light, soil pH, and bare soil (see Table 1) improved the model fit. If so, we kept the squared term of the variable in question instead of the linear effect. After the initial fit and checking for non-linearity as described above, we ran a backward model selection procedure for each model based on AIC. The procedure stopped when AIC did not drop anymore and ΔAIC was above 2 (Burnham and Anderson 2002). In each iteration, we dropped the variable causing the smallest change in AIC value. As a final step, we checked model residuals to ensure that these were normally distributed. We did not conduct spatial modelling in this study, since the original field-work design lying behind the data was designed to avoid spatial autocorrelation, and tests for all the species groups originally inventoried concluded that spatial signals were of minor to little importance (see Brunbjerg *et al.* (2019), and also “Study area and site selection”).

## Results

The two relative fungal dark diversity estimates (based on fungi-only and both fungi- and plant co-occurrences) were between 0.17 – 0.93 (open habitats, median 0.51) and 0.20 – 0.63 (woodland, median: 0.39); and 0.21 – 0.95 (open habitats, median: 0.56) and 0.24 – 0.7 (woodland, median: 0.44) respectively. In most cases, our models explained between 20-30 % of the variation in fungal dark diversity and more than 40 % for the woodlands models when including both LiDAR and measured variables (Table 2). In the “fungi-only dark diversity” model for woodland habitats (Table 2) the squared term of soil pH did improve model fit, but this term was left out during the subsequent model selection procedure.

**Table 2.** Modelling coefficients for the best models (i.e. after model selection) regressing dark diversity estimates based on only fungi co-occurrences (*Fungi-only dark diversity*) or based on both fungi and plant co-occurrences (*Fungi-plant dark diversity*) against the selected explanatory variables. Significant variables were from either a LiDAR-only or a full model with both LiDAR-based and field-measured predictors.

|  |  |  | R<sup>2</sup> | LiDAR variables |  | Field-measured variables |  |  |
| --- | --- | --- | --- | --- | --- | --- | --- | --- |
|  |  |  |  | Amplitude | Echo ratio | Plant richness | Soil fertility | Soil moisture |
| Woodland habitats | Fungi-only dark diversity | Lidar-only model | 0.24 | 0.03 | - | - | - | - |
|  |  | Full model | 0.40 | 0.04 | - | -0.05 | - | - |
|  | Fungi-plant dark diversity | Lidar-only model | 0.23 | 0.03 | - | - | - | - |
|  |  | Full model | 0.44 | 0.04 | - | -0.05 | - | - |
| Open habitats | Fungi-only dark diversity | Lidar-only model | 0.10 | -0.05 | 0.01 | - | - | - |
|  |  | Full model | 0.30 | - | 0.01 | -0.08 | 0.06 | 0.04 |
|  | Fungi-plant dark diversity | Lidar-only model | 0.07 | -0.04 | 0.01 | - | - | - |
|  |  | Full model | 0.34 | - | 0.01 | -0.10 | 0.06 | - |

**Appendix 2.**
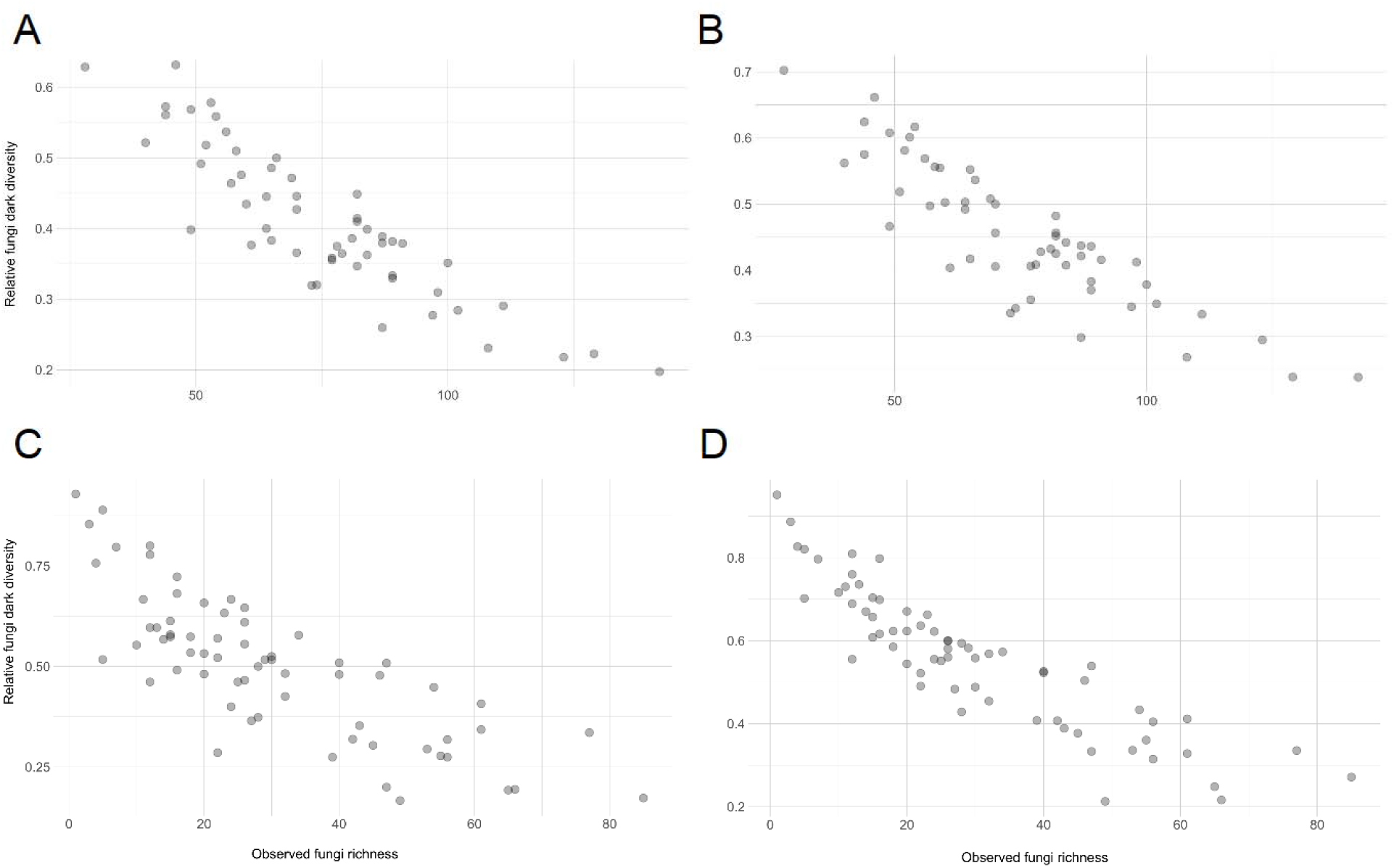
Relationship between observed fungal diversity and dark diversity estimates in woodland based on (A) only fungi co-occurrences (*Fungi-only dark diversity*) or (B) fungi and plant co-occurrences (*Fungi-plant dark diversity*), and in open habitats based on (C) only fungi co-occurrences (*Fungi-only dark diversity*) or (D) fungi and plant co-occurrences (*Fungi-plant dark diversity*).

The only LiDAR variable significant in both open habitats and woodlands was amplitude, which was significant in all models for woodlands and in LiDAR-only models for open habitats (Table 2). This variable had a positive effect on dark diversity in woodlands (Table 2) but a negative influence in open habitats (Table 2). Echo ratio was the only other significant LiDAR variable in our analyses and positively influenced dark diversity in open habitats (Table 2). Plant richness was negatively related to local fungal dark diversity and had the strongest impact of all the field-measured factors included in our analyses (Table 2). Also, soil fertility and moisture were positively correlated with fungal dark diversity in open habitats (Table 2).

In all cases, models considering only the structural environment (LiDAR only) were outperformed by models considering plant richness and the abiotic environment in addition to the structural, notably in the open habitats (Table 2). Appendix 1 shows all pair-wise Spearman correlations between the explanatory variables used and Appendix 2 shows observed fungi richness against the two dark diversity measures for both open and woodland habitats.

## Discussion

In this study, we demonstrate for the first time that LiDAR derived variables, alone and in combination with field-measured variables, can explain a significant amount of the variation in local dark diversity of temperate macro-fungal communities. Our findings indicate that the dark diversity of fungi is influenced by the local vegetation structure, plant associations, and the abiotic environment. This is not surprising since local observed fungal diversity is also determined by these factors to a large degree (Thers *et al.* 2017, Yang *et al.* 2017, Moeslund *et al.* 2019). Indeed, the relative dark diversity analyzed in this study correlated with the observed fungal species richness (Appendix 2), which was expected given that the dark diversity calculations are based on the species present (Noreika *et al.* 2020). On the other hand, there are differences between these two measures, so the results presented here are not necessarily applicable to the observed fungal diversity. We also find that models including field-based variables explained the dark diversity of fungi far better than models relying solely on LiDAR, notably in open landscapes. While LiDAR has the advantage that one can record data from huge areas in very fine detail for relatively low cost, our results indicates that to get the best explanation of local fungal diversity patterns fieldwork is still needed.

### LiDAR-based measures

This study shows that LiDAR captures habitat characteristics potentially important for fungal dark diversity which are not represented by traditional field-measured variables. Notably, the relationship between fungal dark diversity and LiDAR-derived vegetation structure in woodlands was relatively strong. Although LiDAR can successfully quantify biophysical characteristics in all types of habitats, it is known to be more effective in forested habitats (Su and Bork 2007), supporting these findings. The most important LiDAR variables for modelling fungal dark diversity were amplitude and echo ratio which gives us important insights into what environmental aspects, which are not typically recorded in field-surveys, can be captured by a LiDAR approach.

The LiDAR measure of amplitude is sensitive both to surface reflectivity and to the number of targets hit by the laser pulse (Moeslund *et al.* 2019). The lower the reflectivity and the more targets between which the light energy is distributed, the lower the amplitude associated with a given point. This would result in high amplitudes in flatter surfaces while yielding low amplitudes in tall and more complex canopies, or translucent surfaces such as leaves. Hence, this variable can be a proxy for succession stage, surface evenness, or vegetation density; since both flat and sparsely vegetated as well as densely vegetated canopies preventing light penetration will yield high amplitude. Supporting this, amplitude was positively correlated with vegetation height and vegetation cover (denser vegetation resulted in higher amplitude) and negatively correlated with echo ratio (vegetation complexity, see below) and canopy openness. In woodlands, the dark diversity of fungi was positively related to amplitude, suggesting that more species are missing in the relatively tall and dense forests compared to more complex woodlands (e.g., with canopy openings). The positive association between LiDAR amplitude and dark diversity could therefore be a consequence of communities in older well-developed shrubland or old-growth forests with windthrows or other openings, having allowed fungi more time to become established with their associated plants (Fernández-Toirán *et al.* 2006, Twieg *et al.* 2007). Indeed, among the top half of woodland plots with regards to amplitude were plantations (which often have dense canopy and equal-aged trees) and most of them contained high relative fungal dark diversity, while the bottom half of the plots, those having the lowest amplitude and dark diversity, were mostly old forests or shrublands with a more well-developed vegetation structure (e.g., dead or fallen wood, or complex sub-canopy layer).

In open habitats, amplitude and echo-ratio were negatively and positively related to fungal dark diversity, respectively. These results indicate that fewer species are missing from the more even early-successional grasslands without trees and shrubs. We suggest this could be the result of encroachment due to the widespread abandonment of ancient grassland management practices resulting in a loss of small-statured typical grassland plant species without a corresponding gain in species associated with scrub and woodland. It could also reflect that fewer species are missing from calcareous or sandy grasslands since open limestone and white sand have a relatively high reflectivity.

### Plant richness

The most important field-measured variable for modelling fungal dark diversity was plant species richness which was negatively related to dark diversity in both open habitats and woodlands. Plant richness and composition are well-known to correlate with fungal richness and composition (Zak *et al.* 2003, Chen *et al.* 2017, Yang *et al.* 2017, Brunbjerg *et al.* 2018, Wang *et al.* 2018), and sites with lower plant species richness have previously been found to have a relatively higher proportion of plants in the dark diversity (Fløjgaard *et al.* 2020). Here, the negative relation between plant richness and fungal dark diversity may be attributed to greater plant richness frequently associated with more stable communities and ecosystems (Kuiters 2013, Pellkofer *et al.* 2016, Yang *et al.* 2018), which could indicate longer continuity and hence time for fungi to establish. Alternatively, host specific fungi species could be missing due to absence of their symbiotic plant species (Dickie 2007). However, in our study, plant richness had almost the same effect on both fungal dark diversity accounting for present plant species and where these presences where unaccounted for. This points to plant hosts playing a minor role for fungal dark diversity, which is unsurprising as it likely indicates that the calculation of fungal dark diversity using the Beals’ index does not allow fungi having non-present hosts into the site-specific species pool. Another possible explanation is that plant richness mirrors human impact. Generally plant species richness have declined over several decades and continue to as a consequence of agricultural intensification and abandonment of extensive land-use (Hülber *et al.* 2017). Other studies have found human disturbance to be strongly related to fungal richness and dark diversity patterns (Epp Schmidt *et al.* 2017, Pärtel *et al.* 2017a), and future studies may help to tease apart these effects.

### Abiotic environment

Soil fertility is often associated with fungal diversity (Balser *et al.* 2005, Kalliokoski *et al.* 2010, Sterkenburg *et al.* 2015) and was found to have a positive relationship with dark diversity in open habitats. In general, soil fertility influences plant species richness negatively through asymmetric competition (Buckland and Grime 2000, Dybzinski *et al.* 2008, Nadeau and Sullivan 2015, Luo *et al.* 2017). This could explain the negative relationship between the local dark diversity of fungi and soil fertility: lower plant species richness yields a higher dark diversity (see discussion above on “Plant richness”). However, the effect might also be uncoupled from plants and simply due to changes in the soil decomposition microbiota from fungal to bacterial dominance along a gradient of soil fertility and pH (Blagodatskaya and Anderson 1998). Another alternative explanation is that soil fertility affects the density of soil mycophagous and microarthropod species (Cole *et al.* 2005) which also may affect fungal dark diversity (Crowther *et al.* 2013). However, while this explanation might be plausible, the underlying mechanisms are largely unknown, calling for further research to dissect the interactions between soil fertility, soil microarthropods and fungal diversity.

We also found soil moisture had a positive relationship with fungal dark diversity in open habitats. Moisture is known to influence fungal communities (Gómez-Hernández and Williams-Linera 2011, Gupta *et al.* 2018, Frøslev *et al.* 2019) as it affects the growth, colonization rate, and spore and fruit body production (Salusso and Moraña 1995, Jacobson 1997, Kennedy and Peay 2007). The relationship between fungi and plants is probably important in this regard because soil moisture causes a high turnover in plant species composition (e.g., Moeslund *et al.* (2013), and in turn, affects the quality and availability of resources for below-ground fungal communities (Chen *et al.* 2017). Additionally, high soil moisture is a strong environmental filter excluding most macro-fungi species from the wet habitats (Heilmann-Clausen *et al.* 2019). This filter may also be the main reason for the higher fungal dark diversity found in the wet habitats when plant co-occurrences was not included in determining the site-specific species pool. On the other hand, the interdependencies between fungi and plants, and the strong link between plant communities and soil moisture gradients (Xiong *et al.* 2003, Silvertown *et al.* 2015, Valdez *et al.* 2019), may explain why soil moisture was not significant in models where plant co-occurrences was included in determining the site-specific species pool, as this approach perhaps accounts for these interactions.

## Uncertainties

One drawback of basing a study on an organism group like fungi, which live a mostly hidden life, is the unavoidable issues concerning overlooked species and hidden diversity in general (Milberg *et al.* 2008, Abrego *et al.* 2016). If such errors are biased towards specific plots our results could be affected. However, the field-work behind the dataset used here was planned to avoid this exact issue by including several visits at each site at different times of the year, and we do not believe this confounds our findings. Nevertheless, there is always uncertainty when estimating unknowns, such as which species are actually absent despite the fact that they could indeed be in a given site. In this study we used on of the most comprehensive and best possible data set along with state-of-the-art methods to calculate these estimates (see e.g. (Lewis *et al.* 2016, Moeslund *et al.* 2017), and we therefore believe our results are indeed sound and realistic despite this uncertainty.

## Conclusion

This is the first study to investigate potential drivers of the local dark diversity of fungi using both LiDAR derived vegetation and terrain structure as well as field-measured variables. We showed that local fungal dark diversity is strongly dependent on the environment with vegetation structure, plant diversity, and abiotic factors playing important roles. Also, to our knowledge, this is the first study using cross-species group co-occurrence data to determine species pools. This may be a more ecologically sound methodology than using only one taxon group, especially for interdependent taxonomic groups. Future studies and novel approaches will be required to unravel the causal links between fungal communities and habitat and vegetation characteristics; and to gain a better understanding of how LIDAR-based measures can be interpreted as measures of vegetation and terrain structure. Using LiDAR as a tool to determine dark diversity, in conjunction with ecological field measurements, may provide a valuable tool to better guide conservation and restoration planning by identifying sites with high restoration potential (high dark diversity) and high priority for conservation (low dark diversity, or sites where the fungal communities are more “complete”).

## Acknowledgements

We thank Thomas Læssøe for collecting and identifying macro-fungi. We sincerely thank Aage V. Jensen Nature Fund for financial support to CF, AKB, JM, LD, KC and JV through the project “Dark Diversity in Nature Management”. The Biowide project and REJ was supported by a grant from the Villum Foundation (VKR-023343). MP has been supported by the Estonian Ministry of Education and Research (IUT20–29), and the European Regional Development Fund (Centre of Excellence EcolChange). The authors declare no conflict of interest.

